# Mapping phylogenetic trees to reveal distinct patterns of evolution

**DOI:** 10.1101/026641

**Authors:** Michelle Kendall, Caroline Colijn

## Abstract

Evolutionary relationships are described by phylogenetic trees, but a central barrier in many fields is the difficulty of interpreting data containing conflicting phylogenetic signals. Obtaining credible trees that capture the relationships present in complex data is one of the fundamental challenges in evolution today. We present a way to map trees which extracts distinct alternative evolutionary relationships embedded in data and resolves phylogenetic uncertainty. Our method reveals a remarkably distinct phylogenetic signature in the VP30 gene of Ebolavirus, indicating possible recombination with significant implications for vaccine development. Moving to higher organisms, we use our approach to detect alternative histories of the evolution of anole lizards. Our approach has the capacity to resolve key areas of uncertainty in the study of evolution, and to broaden the credibility and appeal of phylogenetic methods.

---

A fundamental challenge in the study of evolution is that for a given set of organisms, markedly different phylogenetic trees can be inferred from each combination of input data, software and settings [1, 2, 3]. Reasons for this include lack of informative data, conflicting signals from descent and selection (convergent evolution), and the fact that evolution is not always tree-like: species trees differ from gene trees, and many organisms exchange genes through horizontal gene transfer. Bayesian Markov Chain Monte Carlo (MCMC) inference methods (e.g. BEAST [4] and MrBayes [5]) produce large posterior collections of trees which are hard to summarize and often include considerable tree uncertainty. We present a method to extract sets of complex evolutionary relationships embedded in data. At the heart of our approach is a way to compare the many possible trees reflecting these relationships.

Direct qualitative comparison of trees (for example, visualization with tanglegrams or DensiTree [6] plots) is illustrative but becomes unwieldy and uninformative when the trees are large or differ significantly. Current quantitative tree comparisons suffer from drawbacks including counter-intuitive behavior [7]. For example, the widely-used Robinson-Foulds metric [8] (RF) is hampered by the fact that large distances between trees do not imply large differences among the shared ancestry of most tips [9].

A *metric* is a mathematical notion of distance; specifying a metric gives structure and shape to a set of objects. Each metric on a set of trees defines a *tree space*. The size and complexity of tree spaces present serious challenges; there are (2*k -* 3)!! possible topologies for trees with *k* labeled tips [10]. As an illustration, this means there are 34,459,425 trees with just 10 tips, and 1076 trees with 50 tips.

We present a method to organize groups of trees, capturing and highlighting their differences according to biological significance. Central to our approach is a tree metric [11] which lends itself to clear visualizations of tree space in low dimensions. It enables straight-forward detection of any distinct groups of similar trees. Most analyses rely on a single tree such as the maximum clade credibility tree (MCC) to summarize a posterior collection of trees (more details in Supplement). However, one of the many drawbacks [12] of this is that crucial information can be lost. Our method provides a natural solution to the problem of summarizing complex tree spaces, producing a small number of representative trees that capture the distinct patterns of evolution reflected in the data.

Our metric works by comparing the placement of the most recent common ancestor (MRCA) of each pair of tips in two trees. The trees can be compared according to their topology alone (disregarding branch lengths), or branch lengths can be included to a chosen extent. Full details are given in [11]. Once the distances between a set of trees have been established, they can be ‘mapped’ into a two-or three-dimensional visualization, as we will demonstrate below. This procedure, using multidimensional scaling (MDS) [13], is explained in more detail in the Supplement.

In the evolution and ecology of higher organisms, phylogenetic trees are used to uncover the origins and adaptations of existing species. This is greatly hindered by the difficulty in resolving species trees using nuclear DNA from different loci and/or mitochondrial DNA. Loci may not contain sufficient variation to estimate trees, and often result in discordant trees. Anole lizards, in particular, are a model system for ecological phenomena including reproductive character displacement, adaptation, behavior and speciation [14]. Recently, the first comprehensive phylogenetic analysis of the *distichus* group of trunk ecomorph anoles (based on mitochondrial and nuclear DNA) [14, 15] found two main areas of uncertainty in the species tree, reflected in clades with low support values. The first is a group containing anoles from the Bahamas and the North Paleo-island of Hispaniola (Figure 3A, clade with 0.64 support). The other is its sister clade, containing mainly anoles from the South Paleo-island, together with *dominicensis1* from northern Haiti.

To explore this uncertainty, we computed pairwise tree distances according to our metric for a random sample of 1000 posterior trees (Figure 3C). There are eight distinct groups of trees (identified through *k*-means clustering), with the MCC tree (Figure 3A) in the center of the largest one. Most groups are arranged in visually separated clusters, particularly when shown in 3D (Figure S4), and have comparable likelihoods in the posterior. The MCC tree alone cannot capture the distinct, likely patterns of evolution supported by the data. We therefore computed an individual MCC tree for each cluster. These illustrate the clusters’ alternative arrangements of the taxa. The support values for clades in cluster-specific MCC trees are high: the clusters resolve uncertain clades into high-certainty clades. The key differences amongst the clusters are in the placement of *dominicensis1*, which is either sister to south Paleo-island anoles or grouped with anoles from Haiti and the Bahamas. This affects the likely origins of the Bahamian anoles, which are grouped along with *dominicensis1* but are closer to either North or South Paleo-island anoles in tree groups of equal likelihood. Our approach has uncovered distinct well-supported patterns of evolution which were not separable by existing methods. The approach obtains analogous distinct patterns using species trees from chorus frogs (see Supplement).

Comparing trees estimated from different genes or loci can play a role in detecting evolutionary behaviors such as horizontal movement of genes and convergent evolution. We compared trees from the 7 genes of Ebolavirus which causes the fatal Ebola hemorrhagic fever. We selected 20 published sequences [16] which differed on every gene (including all five viral species of the Ebolavirus genus) and inferred trees in BEAST from each gene separately and from all genes together. Our method shows (Figure 4) that the VP30 gene, an essential viral transcription activator, has a distinctive phylogenetic pattern, forming three distinct clusters with comparable likelihoods (Figure 4B). Each cluster resolves the uncertainty in the placement of the Sudan clade (pink), placing it in one of 3 positions: sister to the EBOV, TAF and BDBV clade, sister to a larger clade containing all the other sequences, or sister to the Reston clade. The latter placement, which is also found in all the MCC trees other than VP30, agrees with previous analysis [16, 17]. Each cluster places the Reston and Sudan sequences into monophyletic clades, but there are differences in the placement of the Reston 1990 and Sudan 2011 tips.

**Figure 1:**
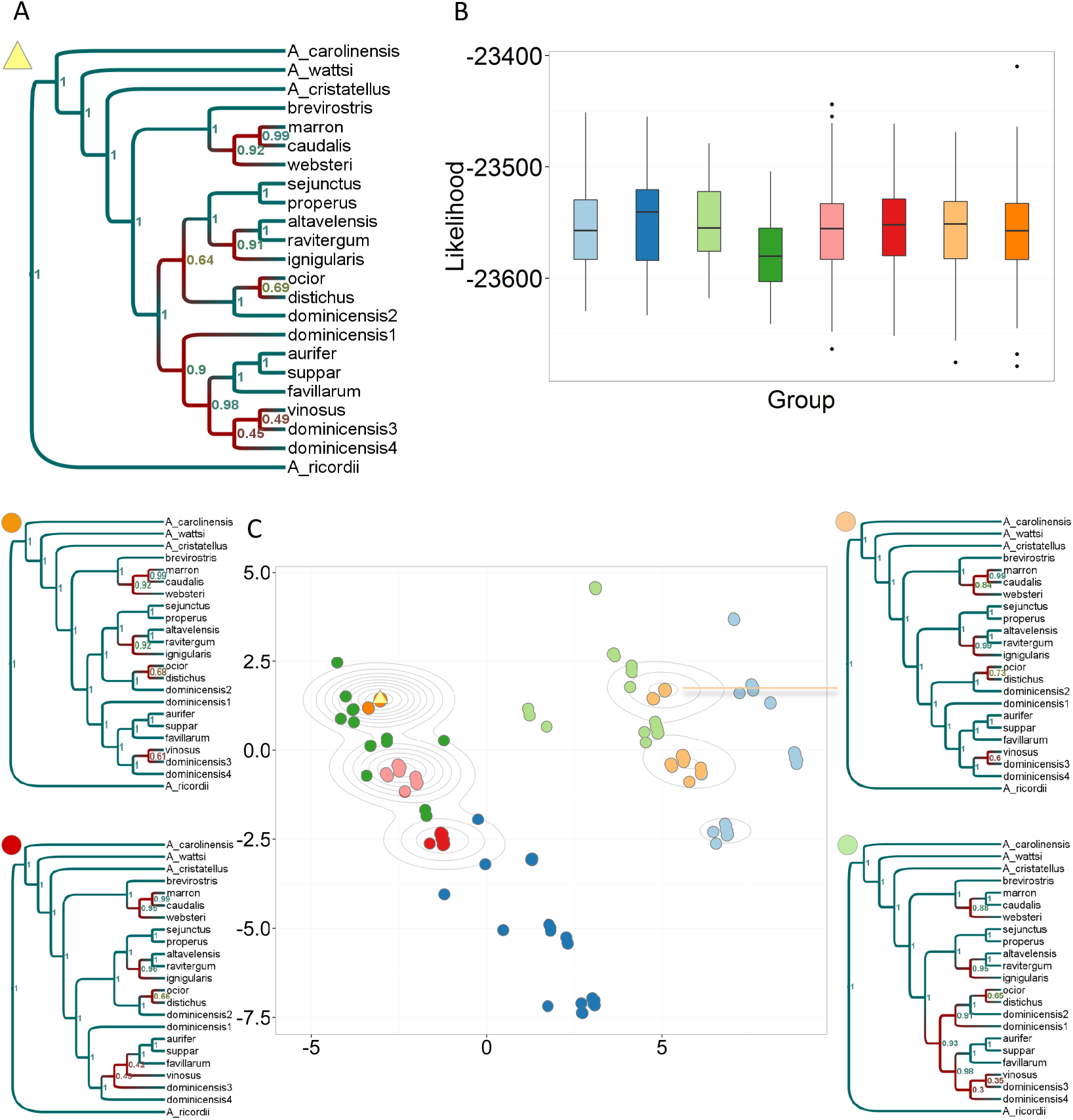
Identifying and exploring islands in anoles *BEAST trees. A: MCC tree for the whole posterior. B: Likelihood distributions were similar for each cluster. C: MDS plot of 1000 trees from the posterior, colored according to 8 clusters found using *k*-means clustering. Four examples of an MCC tree for an individual cluster are given here. A more visually dispersed cluster corresponds to more uncertainty in larger clades. The relatively small number of points in the plot («1000) corresponds to the small number of distinct topologies explored; density of points is illustrated with contour lines.

The alternative phylogenetic signals from VP30, with differences in the deep structure of the tree, suggest a historical recombination event or convergent evolution in this gene. Marzi et al. [18] recently found that a new whole-cell vaccine, EBOVΔVP30 [19, 18], is safe and effective against lethal Ebola challenge in non-human primates. They did not observe recombination in relatively short-term experiments (based on viral passage). However, if VP30 is amenable even to rare recombination events in this genus, this could threaten the future efficacy of EBOVΔVP30 by allowing Ebolavirus to generate vaccine escape strains.

Our method reveals distinct patterns of evolution, both in viruses and in higher organisms. The approach allows quantitative comparisons between the evolutionary patterns of different genes or loci. It can also measure the extent to which a particular locus shapes the most likely phylogenetic trees for a set of data, and thereby identify phylogenetically informative sites, loci or genes. It provides a framework for comparing and testing tree estimation methods themselves [11]. Addressing tree estimation challenges is increasingly important as datasets grow to tens of thousands of tips, rendering standard inference methods infeasible. More generally, our method is relevant to any rooted, labeled trees with the same set of tips, including decision trees, network spanning trees, hierarchical clustering trees and language trees.

**Figure 2:**
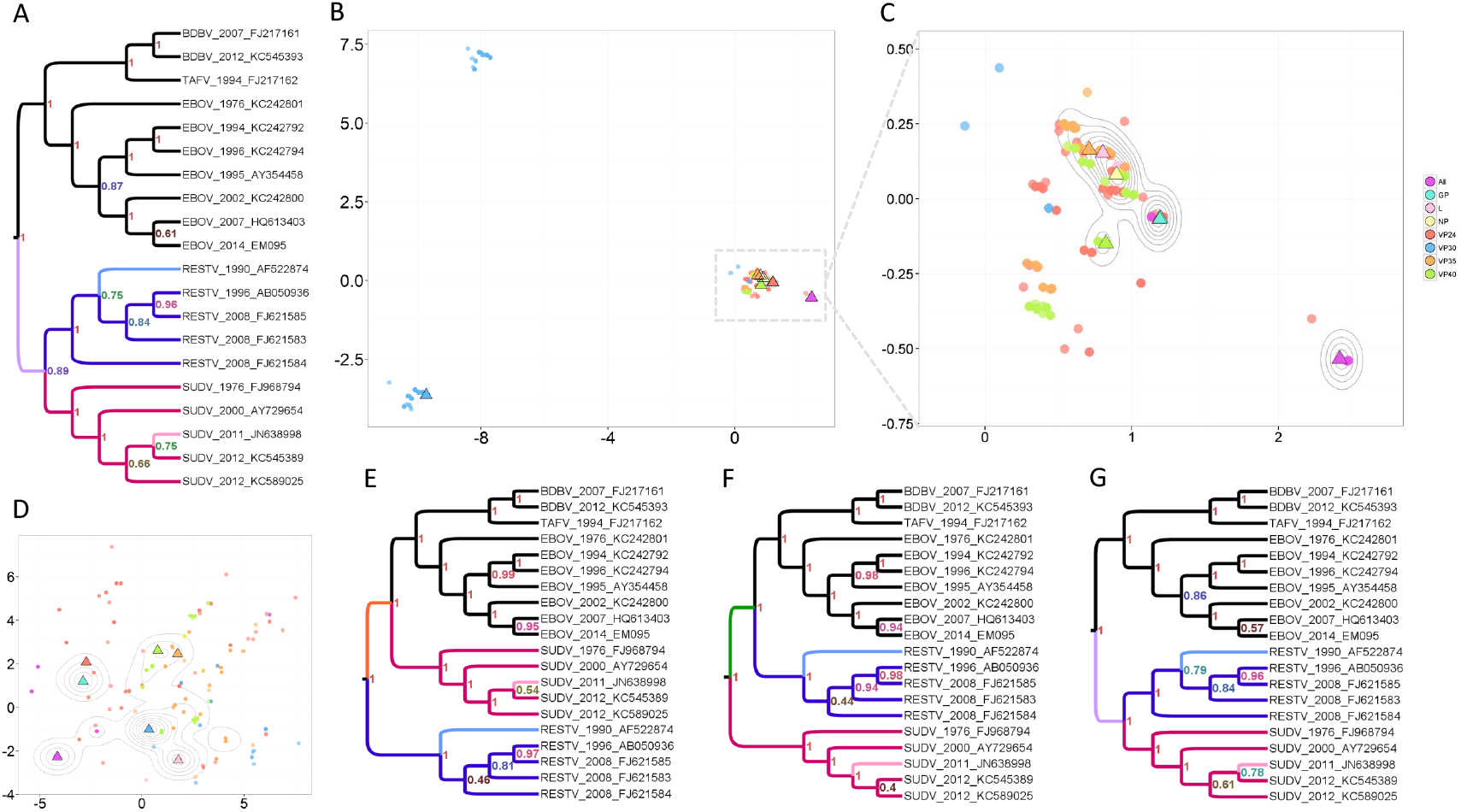
Ebolavirus comparison of individual and ‘all’ gene trees. A: MCC tree from ‘all’ (purple triangle). B: MDS plot of 1200 trees (150 per type), showing three distinct groups of topologies for VP30. C: A closer look at the largest group from B. The MCC tree per gene is marked as a triangle. The MCC trees for GP and VP24 are plotted in almost the same position, in the center of the largest group amongst individual gene trees. The distance between them is 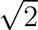, the minimum distance by our metric. D: MDS plot of all 1200 trees using the RF metric. Distinct VP30 topologies are not detected, in fact, the VP30 MCC tree is identical to the NP MCC tree according to RF because it is an unrooted metric. E–G: MCC tree from lower left VP30 cluster, upper left VP30 cluster, and main cluster respectively. The MCC tree from the largest cluster, G, is naturally more similar to A than to E or F.

